# Role of transposons in the specialization of *Botrytis* cinerea to grapevine: Insights into small RNAs and a new *Starship*

**DOI:** 10.64898/2026.05.30.728154

**Authors:** Antoine Porquier, Adeline Simon, Justine Vergne, Jérémy Villette, Sébastien Aimé, Stéphane Bourque, Jérôme Chapuis, Anthony Colas, Justine Rouffet, Antoine Davière, Anne-Sophie Walker, Marielle Adrian, Benoit Poinssot, Muriel Viaud

## Abstract

While transposable elements (TEs) are recognized as major drivers of fungal genome structure, evidence of their direct involvement in the interaction with host plants and the environment is only beginning to emerge. Retrotransposons can generate small RNA (sRNA) that act through cross-kingdom RNA interference, while giant DNA TEs called *Starships* carry dozens of cargo genes that enrich the accessory gene compartment of fungal genomes. In the polyphagous pathogen *Botrytis cinerea,* the Vv3 strain and other strains specialized on grapevine display a specific repertoire of TEs, including the retrotransposons *BcCopia4, BcGypsy6*, and *BcGypsy7*. This study first explored the putative role of sRNA generated from these retrotransposons in the interaction between the Vv3 strain and its host of origin, grapevine. Putative targets were identified among the host mRNAs, but predicted cleavage sites could not be experimentally validated. Moreover, Dicer mutants unable to produce retrotransposons-derived sRNA remained fully pathogenic on grapevine, indicating that these sRNAs do not act as virulence factors on this host. In parallel, this study provides an updated RNA-seq-based annotation of the accessory genes of the Vv3 strain, which revealed a new 93 kb-*Starship* harboring 43 cargo genes, some of which are related to arsenic resistance. A formal genetic approach confirmed that this locus confers resistance to this metalloid. This giant TE, named *Ariane,* was also detected in additional grapevine-specialized strains resistant to arsenic but not in strains isolated from other hosts such as tomato. In conclusion, this study highlights how a *Starship* giant transposon shaped the accessory genes compartment of the polyphagous fungus *B. cinerea* and may have contributed to its adaptation to vine cultivation by conferring resistance to arsenic, a compound widely used in vineyards during the last century.

**IMPACT STATEMENT:** Fungal genomes contain many families of transposons whose functional role in adaptation to the environment and in biotic interactions remained hidden for a long time. In the grey mold fungus *Botrytis cinerea*, strains specialized on grapevine, such as Vv3, carry a specific repertoire of transposons which provides a valuable opportunity to investigate their role in niche adaptation. In this study, we first investigated retrotransposon-derived small RNA, previously described as effectors capable of manipulating the immunity of the model plant *Arabidopsis thaliana*. Although *in silico* analysis of the specific repertoire of small RNAs of the Vv3 strain suggested that some could target the expression of grapevine genes, a genetic approach demonstrated that they do not play a significant role in virulence on this host. In contrast, this study identified a new transposon, named *Ariane,* that carries 43 cargo genes and confers a selective advantage to the Vv3 strain. *Ariane* belongs to a family of giant transposons called *Starships*, recently discovered in fungi and considered to be responsible for horizontal genes transfers between unrelated species. *Ariane* was detected only in some *B. cinerea* strains isolated from grapevine, and a genetic cross showed that it provides these strains with the ability to grow in presence of arsenic. Arsenic was used in vineyards until the beginning of the 21^st^ century to control fungal trunk diseases and insect pests. Therefore, *Ariane* appears to have played an important role in the adaptation of *B. cinerea* strains to cultivated grapevine. Overall, these results underline the importance of considering *Starships* when predicting emergence of resistance to antifungal compounds.

**DATA SUMMARY:** The novel data described in this study, *i.e.,* RNA-Seq data and the *Starship* element are accessible under NCBI GEO accession GSE327899 and at https://doi.org/10.57745/HYWRNM, respectively. All information related to *Botrytis cinerea* genomes used in this study are centralized and kept up to date at the Bioinfo Bioger genomic web portal: https://bioinfo.bioger.inrae.fr/portal/genome-portal/. Direct links to individual portals are respectively https://bioinfo.bioger.inrae.fr/portal/genome-portal/3/ for *B. cinerea* Vv3 genome, https://bioinfo.bioger.inrae.fr/portal/genome-portal/2/ for *B. cinerea* Sl3 genome, and https://bioinfo.bioger.inrae.fr/portal/genome-portal/4/ for *B. cinerea* populations isolated on tomato or grapevine. Each portal provides: (i) a centralized access to public genomic resources, including the genome, transposon, and RNA repositories; (ii) a data browser to download the genomic files; (iii) a genome browser that enables visualization of features within their genomic context, along with associated expression data. As a summary, the prior main public *B. cinerea* genomic accessions and resources used in this study are: GCA_039644125 for VV3 genome, GCA_022560135 for Sl3 genome, GCA_000143535 for B05.10 genome, PRJNA624742 for populations, https://doi.org/10.57745/HYWRNM for transposons, and GSE181592 for small RNAs. Furthermore, table S1 summarizes the list and characteristics of the 64 *B. cinerea* genomes publicly available to date. The genomic data for *Vitis vinifera* genome PN40024.v4 used in this study are available at: https://integrape.eu/resources/genes-genomes/genome-accessions/.

## INTRODUCTION

Transposable elements (TEs) are recognized as major drivers in the evolution of fungal genomes, owing to move between genomic loci and to favor genomic rearrangements. In fungal plant pathogens, TE-rich compartments provide a favorable environment for the diversification of effector gene repertoires [1]. Insertions of TEs in promotors can also lead to gene overexpression which in turn may confer antimicrobial resistances such as in the wheat pathogen *Zymoseptoria tritici* [2]. In this species, a large-scale population study revealed historic TE mobilization waves that generated distinct pools of adaptive variants [3]. While the functional consequences of TE genomic localization are now well described, direct involvement of both retrotransposons and DNA transposons in host–pathogen interactions through their own sequences also emerges. Retrotransposons have been shown to be sources of small interfering RNAs (siRNAs) that act as effectors in cross-kingdom RNA interference (CK RNAi), allowing fungal pathogens to suppress host immunity [4]. Regarding DNA TEs, two families of large elements able to carry cargo genes were identified. First, *Helitrons* were described for their capacity to capture genes or fragments of genes, which increases their size [5]. More recently, giant TEs called *Starships* that can reach up to 700 kb and carry dozens of cargo genes were discovered [6–8]. Inside a fungal species, these cargo genes enrich the compartment of accessory genes which can provide a selective advantage. Several cargo genes of *Helitrons* and *Starships* were shown to directly shape fitness and virulence-related traits [9–12].

The grey mold fungus *Botrytis cinerea* is notable for its ability to infect a wide variety of hosts [13] and is therefore considered generalist. However, as for some other generalist pathogens, it in reality consists of multiple coexisting populations, each exhibiting a degree of host specialization. Notably, genomic studies revealed that French populations of *B. cinerea* isolated from tomato (Sl strains) and grapevine (Vv strains) correspond to different genetic groups and have higher virulence on their host-of-origin than other strains, indicating host specialization [14, 15]. In a complementary study, the genomes of the representative strains Vv3 and Sl3 were fully assembled using the PacBio technology and this revealed that they differ in their accessory chromosomes (ACs) and accessory genes in subtelomeric regions of core chromosomes as well as in their repertoires of TEs. In particular, the Vv3 and a majority of strains specialized on grapevine carry the TE-rich AC 19 that is absent from Sl3 and other strains specialized on tomato [16]. The identity and density of (TEs) were clearly different between strains, with a larger number of subfamilies (26) and a greater genome coverage in Vv3 (7.7%) than in Sl3 (14 subfamilies, 4.5% coverage). Notably, six retrotransposons of the *Copia* and *Gypsy* families were present in Vv3 but not in Sl3. Three of them (*i.e. BcCopia4, BcGypsy6* and *BcGypsy7*) were identified as templates for the synthesis of small RNAs (21-22 nt). Extending the study to additional Vv and Sl strains indicated that *BcCopia4* and *BcGypsy7* and their derived small RNAs are common features of the main population specialized on grapevine and were not found in strains isolated from tomato [16]. Importantly, it has been shown by Weiberg et al. [4] that *B. cinerea* retrotransposon-derived siRNAs can act as effectors of cross-kingdom RNA interference (CK RNAi). Thanks to this pioneering study and additional ones all conducted with the model strain B05.10, a comprehensive view of the retrotransposon-derived siRNAs synthesis pathway in *B. cinerea* and their action in the host plant is now available. When transcribed, ORFs of *B. cinerea* retrotransposons are used as templates by the RNA-dependent RNA polymerase BcRDR1 to generate double-stranded RNA (dsRNA) precursors [17]. These dsRNAs are then cleaved by the Dicer-like (DCL) proteins BcDCL1 and BcDCL2 to yield 21-22nt long sRNAs. It was demonstrated that during the infection of *A. thaliana*, these fungal sRNA are loaded on the host AGO1 protein and direct the cleavage of target messenger RNAs of genes involved in immune response thereby acting as effectors [4, 18].

The overall objective of this study was to investigate the role of the specific repertoires of TEs and their potential cargo genes in *B. cinerea* strains specialized on grapevine. Considering the role of retrotransposons-derived sRNAs in CK RNAi, we investigated their role in the interaction with grapevine. Grapevine mRNAs potentially targeted by sRNAs derived from *BcCopia4, BcGypsy6*, and *BcGypsy7* TEs were investigated using *in silico* prediction tools, expression analysis and cleavage validation assays. In addition, generation of Dicer mutants in the Vv3 strain allowed to evaluate the contribution of fungal sRNA to virulence toward grapevine. In a second approach, the presence of large DNA TEs and cargo genes among the accessory genes of the Vv3 strain was explored. Based on newly generated transcriptomic data, we annotated a previously undescribed *Starship* element harboring 43 cargo genes. This giant TE named *Ariane* was detected in additional strains specialized on grapevine and was experimentally shown to confer resistance to arsenic, a compound that was widely used in vineyards during the last century.

## METHODS

### *Botrytis cinerea* strains, growth conditions and standard molecular methods

The Vv3 and Sl3 strains were isolated in the Champagne region (France) from grapevine and tomato respectively [15] and their fully assembled genomes were described in Simon et al. [16] (Table S1). A cross between the Vv3 and Sl3 strain was performed as previously described [19] and one hundred progenies were isolated for genetic analyses. Routine cultivation of the different strains was carried out on rich medium (ML; 20 g/L malt extract, 5 g/L yeast extract and 15 g/L agar) at 20°C with an alternation of 12 h of white light and 12 h of darkness. To obtain the formation of sclerotia, complete darkness was used. The growth curves of the different strains were compared by nephelometry [20]. For inoculum preparation, conidia were collected from 10-day-old solid cultures and suspended in Potato Dextrose Broth (PDB, Disco) or minimal medium (MM; 20 g/L sucrose, 1 g/L KH_2_PO_4_, 0.5 g/L KCl, 0.5 g/L MgSO_4_ 7H_2_O, 0.01 g/L FeSO_4_ 7H_2_O, 3 g/L NaNO_3,_ pH 5.0) to obtain a concentration of 10^4^ conidia/mL. For each strain, eight wells of a 96 wells-microplate (Sarstedt) were filled with 300 µL of suspension. Growth was automatically recorded each 15 minutes during 72h using a nephelometric reader (NEPHELOstar Plus, BMG Labtech, Omega 5.70), equipped with a 635-nm laser as radiation source. Measurements were done with a laser beam focus of 2.5 mm and an intensity of 80%. The 96-well plates were shaken at 400 rpm, with a double orbital movement between readings. Fungal DNA and RNA were extracted as previously described [21]. Standard PCR analyses were conducted with the GoTaq polymerase (Sigma). All PCR primers are listed in Table S2.

### Prediction of the grapevine transcripts targeted by fungal small RNA

Libraries of small RNAs of the *B. cinerea* strains Vv3 and Sl3 were previously obtained from mycelium growing on grape juice medium [16]. Small RNAs were filtered to exclude sequences present in less than 50 reads per million (rpm), and collapsed into unireads. Unireads were submitted to the TAPIR tool [22] to identify potential target candidates among *V. vinifera* (PN40024.v4); *i.e.,* the tapir_fasta program was run on 20 to 24 nt unireads with score <= 4.5 and mfe ratio >= 0.7.

### Transcriptomic analysis of infected grape berries

Grape berries (*Vitis vinifera*, cv. Italia) were infected with conidia of *B. cinerea* Vv3 and Sl3 strains as previously described [23]. In short, the equatorial area of each berry was inoculated with 15,000 conidia, in potato dextrose browth (PDB; Duchefa Biochimie) diluted to the fourth. The inoculated berries were incubated at 20° C with 100% relative humidity in a growth chamber with 12 hours of light per day. At 24 and 48 hours post-inoculation (hpi), skin samples (without pulp, *i.e.,* exocarp) of 5 mm diameter were collected at the inoculation sites. Three replicates of 20 berries were performed per modality. Total RNA was extracted with a procedure optimized for grape tissues [23] and submitted to a DNAse treatment (Ambion). The 12 total-RNA samples were used for RNA-SEQ sequencing. Illumina TruSeq Stranded mRNA Sample Prep kit was used to prepare all RNA-seq libraries, following the manufacturer’s instructions. Sequencing was performed by Integragen (https://www.integragen.com/). Samples were run on an Illumina HiSeq 4000 in paired mode, 2×75 bp. About 35 million pairs were sequenced for each sample. Quality of raw sequence data was checked with FastQC [24]. Reads were mapped using STAR [25], against *V. vinifera* genome (PN40024.v4) with parameters ––alignIntronMax 60000 ––alignMatesGapMax 60000, and against the genome of the *B. cinerea* strain Vv3 [23] with parameters ––alignIntronMin 20 ––alignIntronMax 5000 ––alignMatesGapMax 5000. The matrices of *V. vinifera* and *B. cinerea* reads counts were analyzed using SARTools [26] for quality controls and for computing the DESeq2 [27] normalization and differential expression. Read mapping to plant and fungal genomes confirmed sufficient sequencing depth for downstream analyses of both organisms. Transcripts with adjusted p-value <0.05 and basemean > 50 were considered as statistically differential if fold change > 1.5 in *V. vinifera* analysis.

### AGO pull-down

AGO1 pull-down assays were conducted with grapevine cell suspensions and vitroplants produced as previously described [28, 29]. Total proteins were extracted by bead-milling (TissueLyser extraction) in a buffer containing 50 mM HEPES pH 7.5, 5 mM EGTA, 2 mM DTT, and a protease inhibitor cocktail (MedChemExpress). Approximately 300 µg of total proteins were incubated with 50 µL of magnetic beads (Sephira, cat. no. 28951378), together with 1 µL of anti-AGO1 antibody (Agrisera, AS09 527) or an antibody previously validated for AGO1 immunoprecipitation in *Arabidopsis* [30] in a final volume of 1 mL of PBS buffer supplemented with 137 mM NaCl and 2.7 mM KCl. Potentially immunoprecipitated proteins were subsequently analysed by western blotting using anti-AGO1 antibodies.

### RLM-RACE PCR

Total RNAs from infected berries (4 samples for each condition) were extracted as described above. All the extracted RNAs (between 1 to 10 µg for each sample) were used to purify mRNAs using the NEBNext Poly(A) mRNA Magnetic Isolation Module (Qiagen). The 4 samples for each condition were pooled and concentrated using the Monarch® Spin RNA Cleanup Kit (10 μg) from NEB allowing to get around 80 ng of mRNAs. For the RLM-RACE (RNA ligase-mediated 5′ RACE) approach, the protocol proposed by Neller et al. [31] was adapted. Briefly, a 5’ RNA adapter (80 ng) was ligated to the mRNAs pools with exposed 5’ monophosphate. The mRNA pools were then reverse transcribed using a 3’ d(T) adapter. After a first non-specific amplification of the pools of cDNA targets, two successive gene-specific amplifications were performed (Primers in Table S2). The resulting PCR fragments that displayed the expected size on gels were cloned in pUC19 (thanks to overlapping regions in the used primers) and sequenced.

### Inactivation of the Dicer encoding genes *BcDcl1* and *BcDcl2*

The CRISPR/Cas9 procedure developed for *B. cinerea* by Leisen et al. [32] was used to inactivate *Bcdcl1* and/or *Bcdcl2* genes in strain Vv3. Inactivation cassettes were generated by PCR (Q5® High-Fidelity DNA Polymerase, NEB) using primers containing 60 bp tails that were designed for integrating the hygromycin and nourseothricin resistance genes [33] at the *Bcdcl1* and *Bcdcl2* loci respectively (Table S2). Fungal protoplasts were prepared as previously described [34] using Vinotaste® enzymes (Novozymes) and mixed with both the PCR cassette and the Cas9/single-guide RNA ribonucleoprotein (RNP) composed of 2µl of EnGen® Spy Cas9 NLS (NEB) and approximately 2 µg of sgRNAs (generated with the EnGen® sgRNA Synthesis Kit, NEB). Transformants were selected and subcultured on 50 µg/mL hygromycin (Sigma Aldrich) or 35 µg/mL nourseothricin (Jena Bioscience). The targeted insertion and the absence of wild type nuclei were verified by PCR. For the double *ΔΔBcdcl1/dcl2* mutant generation, two independent *ΔBcdcl1* mutants were used as recipient to further inactivate *Bcdcl2*. Two single mutants and two double mutants (*ΔBcdcl1* A and B, *ΔBcdcl2* A and B, *ΔBcdcl1 ΔBcdcl2* A and B) were selected for this study. One mutant of each genotype was further analyzed by full genome sequencing (Illumina NovaSeq X Plus Series, paired mode 2 x 150 bp). Sequencing reads were mapped onto the *B. cinerea* Vv3 genome [23] (GCA_039644125.1) using BWA [35]. Aligned reads were filtered based on quality using SamTools [36] and Picard Tools [37] to remove secondary alignments, reads with a mapping quality <30, and paired reads not at the expected distance. SNP calling was performed with Freebayes [38]. Further filtering was carried out using script VCFFiltering.py (https://urgi.versailles.inra.fr/download/gandalf/VCFtools-1.2.tar.gz). We kept only SNPs supported by more than 90% of aligned reads, detected outside low-complexity regions or transposable elements (as identified in Vv3 genome) and with coverage lower than twice the standard deviation from the mean depth coverage. In parallel, sequencing reads were assembled using SDAdes [39].

### Stem-loop PCR

Total RNAs from *B. cinerea* strains grown 7 days on ML medium were extracted using Trizol. Detection of BcsRNAs was carried out following the stem-loop RT-PCR protocol [40] using 1 μg of total RNAs. The stem-loop RT-PCR products were visualized on 4% agarose gels.

### Pathogenicity assays

For pathogeny tests on grapevine leaves, plants of *V. vinifera* cv. Marselan were grown in a greenhouse as described previously [41]. In brief, they were produced from herbaceous cuttings planted in individual pots and grown at 16/8 h light/dark photoperiod at 20/18°C until they developed six leaves. Forty 1.9 cm-diameter leaf discs were then excised from the second and third youngest fully expanded leaves and placed on moist Whatman paper. In parallel, conidia of *B. cinerea* were harvested from approximately 10-day-old cultures and resuspended in potato dextrose broth (PDB; 6 g/L; Difco) to a final concentration of 5. 10⁴ conidia mL⁻¹. Leaf discs were inoculated on their adaxial surface with a 20 µL droplet of the conidial suspension (i.e. 1000 conidia) and incubated in plastic containers at 100% relative humidity under a 10/14 h light/dark photoperiod at 20/18°C. Disease development was assessed 3 days post-inoculation by measuring lesion diameters using ImageJ software (https://imagej.net/ij/). Other pathogeny assays were conducted with a similar protocol except that conidia were resuspended in PDB diluted to the fourth. Inoculated grapevine berries (*V. vinifera* cv. Italia) and detached leaves from five-week-old French bean plants (*Phaseolus vulgaris* cv. Caruso) were incubated with a photoperiod of 16 h of white light at 22°C and 8 h of darkness at 18°C.

### Prediction of the Vv3 accessory genes and of the *Starship* element

The annotation of the genes of the Vv3 and Sl3 strains was previously conducted by the transfer of the structural annotation of the genes of the model strain B05.10 [42] supplemented by a *de novo* annotation of the Vv3– and Sl3-specific genes [16]. In this study, prediction of Vv3 accessory genes was updated taken advantage of the RNAseq data: Vv3 genome masking was performed with REPEATMASKER [43] to filter out transposons previously identified [16]. Structural annotation was then performed with GENEMARK-ES [44], AUGUSTUS [45], HELIXER [46, 47] and EUGENE [48] with fungal proteins [49] and transcripts from this study as evidence.

Selection of the best gene models was performed with INGENANNOT [50]. The newly predicted accessory genes of Vv3 replace the previously identified specific genes in Simon et al. [16]. The track for the new annotation replaces the previous one and is displayed alongside the track showing genes transferred from the reference annotation of B0510 on the Vv3 genome browser (See data summary). A functional annotation was performed using INTERPROSCAN [51]. The classification of the *Starship* identified on chromosome 3 (Bcin_Vv3_03: 3.353.355 – 3.446.435) was carried out following the guidelines provided in the Starfish documentation [9], i.e. both YR captains were queried against the HMM profiles derived from reference YR sequences (YRsuperfams.p1–512.hmm) using hmmsearch. Vv3 YR captain proteins and 1222 representative Starship YR) were then aligned using MAFFT [52], trimmed using TRIMAL [53] and phylogeny was constructed using IQ-TREE [54]. Trees were displayed with iTOL [55].

### Growth assays to evaluate arsenic resistance

To test the effect of arsenic on *B. cinerea* germination and hyphal growth, conidia were inoculated in PDB supplemented with different concentrations of AsNaO_2_ (0, 250, 500, 750, 1000 μg/mL) and incubated at 12°C. Germination was observed 20 hpi. Finally, a concentration of 750 μg/mL was used to screen for arsenic resistance in WT strains and in the Sl3 X Vv3 progeny. All germination and growth tests were performed twice and >100 conidia were observed for each sample.

## RESULTS

### Identification of grapevine transcripts putatively targeted by small RNAs that are specifically produced by grape-specialized *Botrytis cinerea* strains

Genomic analyses of *B. cinerea* populations previously identified three small RNA producing-retrotransposons *i.e., BcCopia4, BcGypsy6* and *BcGypsy7* that are present in the strains specialized on grapevine including Vv3 but not in the strains specialized on tomato (Sl strains) nor in the B05.10 model strain. As illustrated in the karyoplot of the Vv3 strain (Fig. 1A), these three TEs occur in multiple copies on the core chromosomes but there is a notable enrichment of *BcGypsy7* on the accessory chromosome AC19 (7 copies out of 23), which is specific to grape-specialized strains. To identify grapevine messenger RNAs (mRNA) putatively targeted and silenced by the sRNAs derived from these three TEs, both in silico and transcriptomic analyses were conducted. Small RNAs sequenced from the strain Vv3 [16] were used to screen the PN40024.v4 grapevine transcriptome using the TAPIR1.1 algorithm which is designed to detect sRNA-target duplexes [22]. From this analysis, 699, 112 and 278 grapevine target transcripts were predicted for *BcCopia4, BcGypsy6* and *BcGypsy7* sRNAs, respectively (Table S3). In total, 1,078 grapevine transcripts were predicted to be targeted by one or several sRNAs. These sRNA were mainly 21 or 22 nt long (94%) and all started with an uracil which is a typical feature of the sRNA that can be loaded on the AGO1 protein [56].

**Fig. 1.**
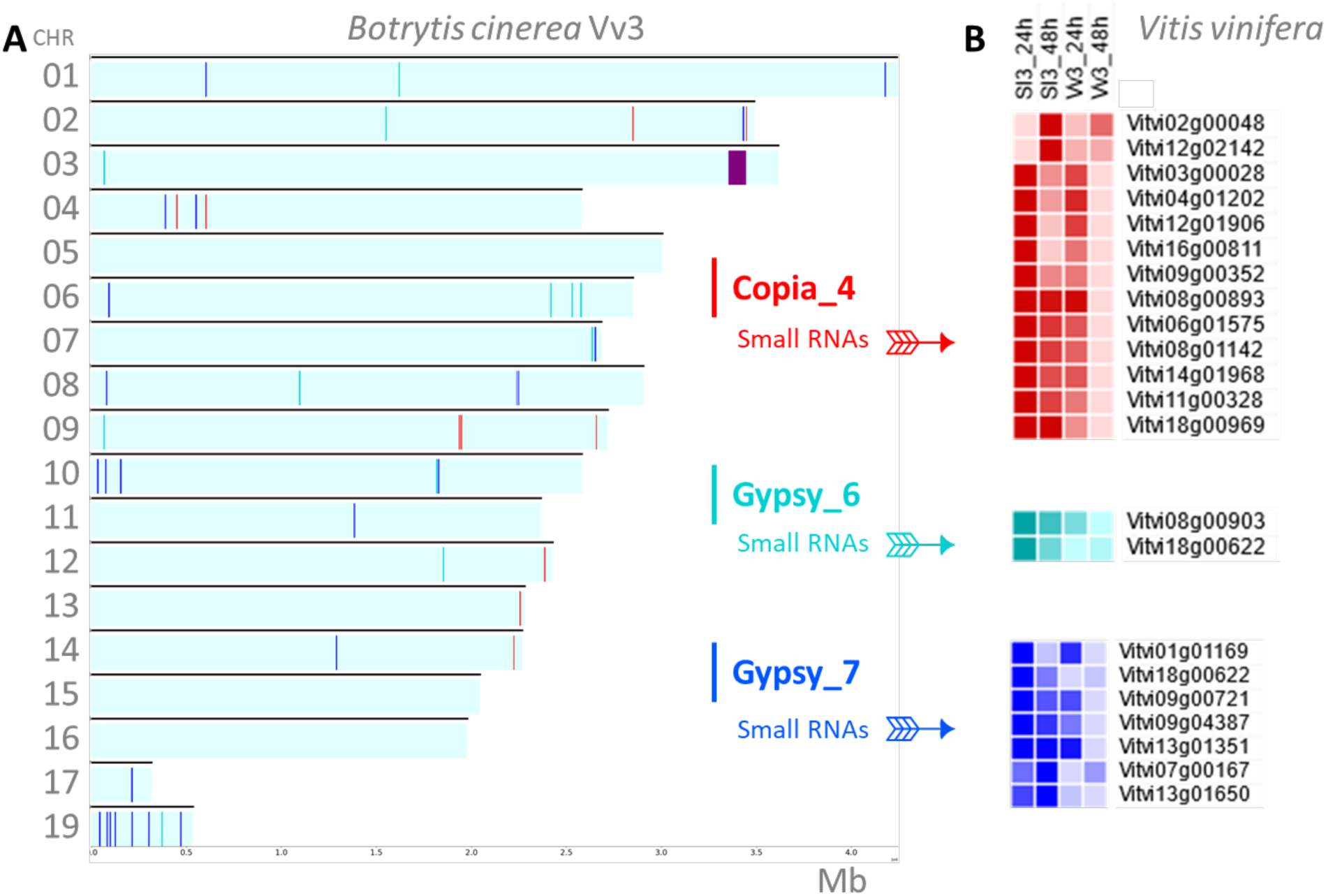
Small RNA-producing retrotransposons *BcCopia4, BcGypsy6* and *BcGypsy7* of the *Botrytis cinerea* strain Vv3 and predicted target genes in grapevine. A. Genomic localization of complete copies of the retrotransposons *BcCopia4, BcGypsy6* and *BcGypsy7,* which are specific to grapevine-specialized strains. The localization of the *Starship* transposon named *Ariane* is also indicated (purple). B. Expression profiles of the 21 grapevine genes predicted to be targets of these retrotransposon-derived sRNAs during infection. These candidate target genes were predicted by *in silico* analysis using the TAPIR1.1 algorithm and were further selected because they showed lower expression levels in berries infected with the Vv3 strain than in berries infected with the Sl3 strain.

If the identified grapevine candidate messenger RNAs are targeted by *BcCopia4, BcGypsy6* and *BcGypsy7* sRNAs during the infection process, their expression is expected to be silenced conversely to what happens when grapevine is infected by a strain that do not possess these retrotransposons *e.g.,* the tomato specialized strain Sl3. Therefore, a RNAseq experiment was designed to compare the transcriptome of grapevine berries infected by Sl3 or Vv3. After 24 or 48 hours post infection (hpi), the berries were scalped to focus on the infection site and total RNAs were extracted to perform RNAseq. In total, 2,423 grapevine genes were differentially expressed between berries infected with Sl3 versus Vv3. In regards with our hypothesis, we focused on genes that were less expressed in berries infected by Vv3 than in berries infected by Sl3 at 24 hpi or 48 hpi. We identified a total of 573 genes following these criteria (Table S4). Out of the 573 corresponding transcripts, 21 were also predicted to be targets of Vv3-specific sRNAs by the TAPIR algorithm, thereby representing possible grapevine candidates for a specific silencing by the Vv3 strain (Table S5; Fig. 1B).

To further investigate whether Vv3 specific sRNA could target the 21 identified candidate grapevine transcripts, we aimed to (i) determine whether the fungal small RNAs were loaded on grapevine AGO1 protein and (ii) validate the cleavage of the 21 grapevine transcripts by the fungal Vv3 sRNAs. The various AGO1 pull-down attempts, aimed at co-precipitating AGO1-bound siRNAs, from undifferentiated grapevine cells or vitroplants, did not allow us to successfully immune-precipitate AGO1 (See Methods). Then, a PCR approach was designed to validate the mRNA cleavage by fungal sRNAs at the sites predicted by the TAPIR1.1 algorithm. Here, we set up a RLM-RACE (RNA ligase-mediated 5′ RACE) approach that takes advantage from the exposed 5’ monophosphate of cleaved mRNAs to highlight the cleavage site [31]. Primers were designed for the 21 candidate mRNAs and the RLM-RACE protocol was applied to RNA extracted from Vv3-infected berries. After two successive specific amplifications, PCR products were sequenced allowing us to monitor the distance between the predicted cleavage site (between the 10^th^ and 11^th^ nucleotide of the sRNA) and the cleavage site revealed by the experiment. The results (Fig. S1) indicated that only one cleavage site was close (1 nt away) from the predicted one (for Vitvi08g00903). Therefore, none of the predicted cleavage sites could be validated by this PCR approach.

### The DICER protein BcDCL1 of the Vv3 strain is required to produce retrotransposon-derived small RNAs

As the predicted cleavage of grapevine mRNAs by fungal siRNA could not be experimentally validated, a reverse genetic approach was conducted to assess the involvement of *B. cinerea* retrotransposons-derived small RNA in its interaction with grapevine. In the *B. cinerea* model strain B05.10, the Dicer-like encoding genes *BcDcl1* and *BcDcl2* were found to have a redundant role in the generation of small RNAs and only the *ΔΔBcdcl1/2* double mutant was impaired in their production [4]. The BcDCL1 and BcDCL2 proteins of the Vv3 strain (thereafter named BcDCL1^Vv3^ and BcDCL2^Vv3^) were found to be 100% identical to the B05.10 proteins. Using the CRISPR-Cas9 procedure [32], knock-out mutants for *BcDcl1^Vv3^*and *BcDcl2^Vv3^* genes as well as double mutants were generated. PCR analyses confirmed the insertion of the hygromycin and nourseothricin resistant cassettes at the targeted loci for six selected mutants *i.e., ΔBcdcl1^Vv3^*A and B, *ΔBcdcl2^Vv3^*A and B, *ΔΔBcdcl1/2^Vv3^* A and B (Fig. S2). For one mutant of each genotype (*ΔBcdcl1^Vv3^*A, *ΔBcdcl2^Vv3^*A and *ΔΔBcdcl1/2^Vv3^*A), the genome was fully sequenced with the Illumina technology and the results confirmed that (i) no additional insertion of the resistance cassette occurred in an ectopic locus, (ii) the mutants were genetically pure (absence of WT nuclei), and (iii) no nonsilent mutations occurred in coding sequences of other genes during the transformation process (Fig. S2).

To assess the ability of the *Δbcdcl1^Vv3^*, *Δbcdcl2^Vv3^* and *ΔΔbcdcl1/2^Vv3^* mutants to produce sRNAs, we conducted a stem-loop RT PCR analysis [40] enabling to highlight the production of specific sRNAs. We focused on the three previously sRNAs characterized in B05.10 *i.e., Bc*siR3.1, *Bc*siR3.2 and *Bc*siR5 [4]. In addition, we designed primers to amplify a Vv3-specific sRNA (*Bc*siR1^Vv3^) predicted to target the grapevine mRNA of the gene *Vitvi02g00048*. As expected from previous sRNA sequencing data [4] [16], *Bc*siR3.1, *Bc*siR3.2 and *Bc*siR5 were detected in the three wild type strains (B05.10, Sl3 and Vv3) while BcsiR1^Vv3^ was detected only in Vv3 (Fig. 2). Analysis of the mutants generated in Vv3 showed that the *ΔBcdcl1* and the *ΔΔBcdcl1-Bcdcl2* mutants are impaired in sRNA production while the *ΔBcdcl2* mutant is still able to produce the three tested sRNA. This suggested that in the Vv3 strain, BcDCL1 is required for the production of retrotransposons-derived sRNAs and that this role could not be taken over by BcDCL2.

**Fig. 2.**
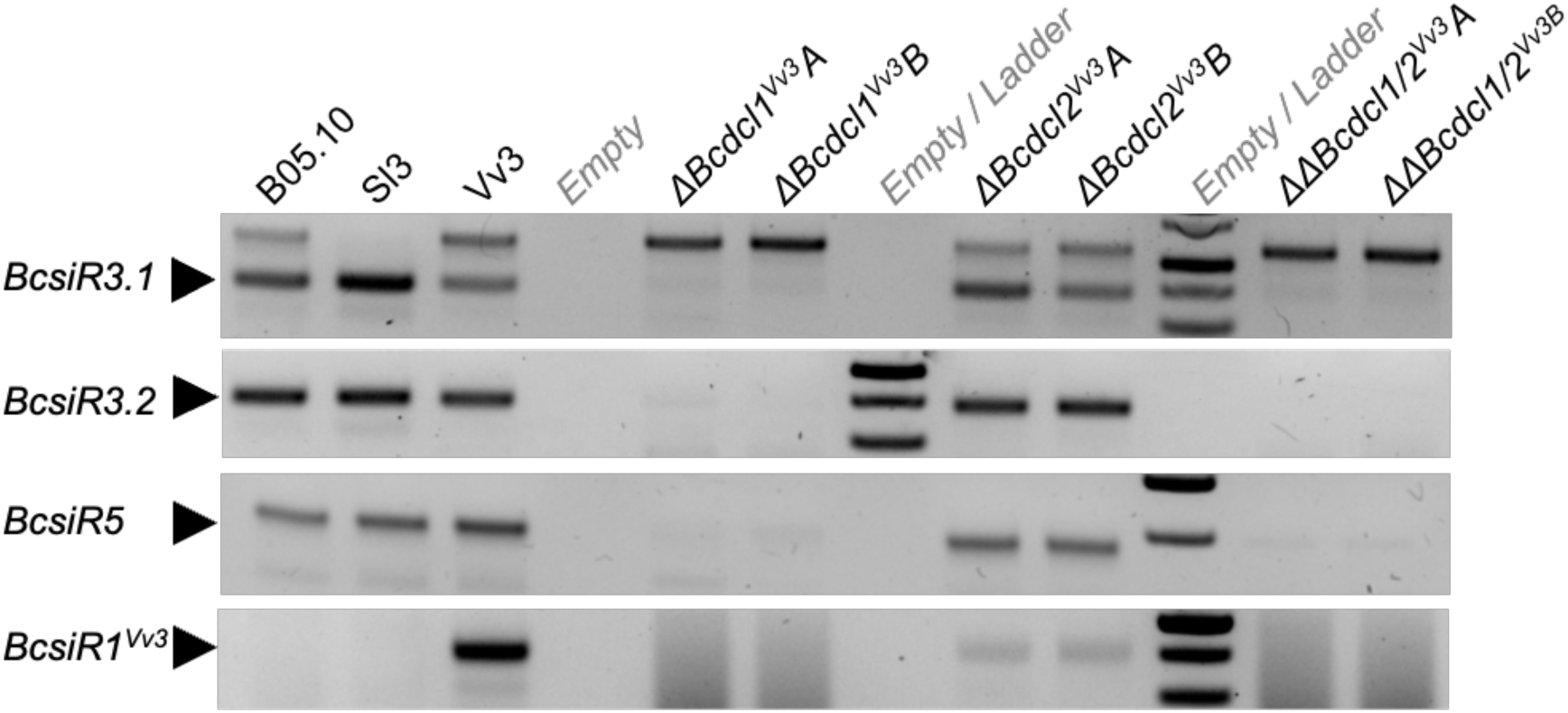
Inactivation of *Bcdcl1* in the *B. cinerea* strain Vv3 impairs the production of small RNAs. Wild-type and Dicer-like mutant strains were grown *in vitro* for seven days on ML medium. Total RNAs were then extracted and the stem-loop RT-PCR was used to detect specific siRNAs. The black arrows indicate the position of the expected band.

### Inability to produce retrotransposon-derived small RNAs does not impact the virulence of the Vv3 strain

On standard cultivation conditions, the different single and double Dicer mutants were not affected in their development and sporulation nor in the production of the survival forms *i.e.,* sclerotia (Fig. S3.). Mycelial development in liquid medium was quantified by nephelometry [20] and the results showed that the WT strain Vv3 and the mutants displayed similar growth curves (Fig. S4). We monitored their virulence on grapevine leaf disks and berries by measuring the necrosis (Fig. 3; Fig. S5). On both organs, the grapevine-specialized strain Vv3 was more virulent than the Sl3 strain, as previously observed [15]. Regarding the different mutants, none of them appeared to be significantly affected in its ability to develop necrotrophic symptoms. Similar results were obtained on *Phaseolus vulgaris* (French bean) detached leaves (Fig. S6). In conclusion, the results suggest that fungal sRNAs do not have a significant role in the virulence of the Vv3 strain.

**Fig. 3.**
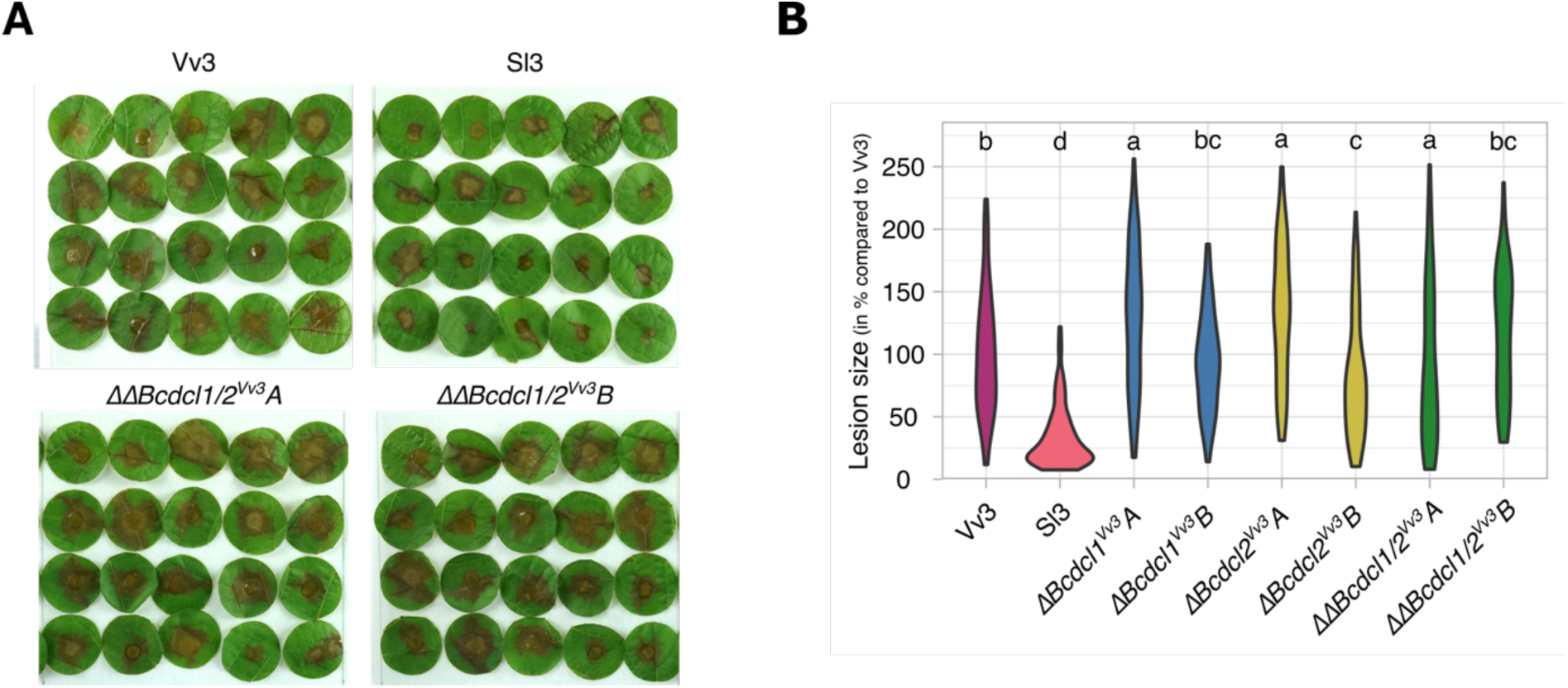
Inactivation of *Bcdcl1^Vv3^*and *Bcdcl2^Vv3^* genes does not impact the virulence of the Vv3 strain on grapevine. A. Leaf disks from *Vitis vinifera* cv. Marselan plants were inoculated with 1,000 conidia of the assessed *B. cinerea* strains (only the pictures for the 2 WT Vv3 and Sl3 and the 2 double mutants *ΔΔBcdcl1/2Vv3*A and B are represented here). The pictures were taken 3 days post-inoculation. B. Necrosis areas were measured based on the pictures using the ImageJ software. The plotted results were obtained from 3 biological replicates (40 disks per assay and per strain representing 120 measures except for *ΔΔBcdcl1/2Vv3B* for which 118 measures were monitored). Statistical groups were defined based on a Kruskal-Wallis analysis.

### Annotation of Vv3 accessory genes reveals a *Starship* transposon of the *Phoenix* family

Beyond the 11,609 genes shared between the Sl3 and Vv3 strains, the Vv3 genome has previously been reported to harbor approximately 200 accessory genes that are absent from the Sl3 genome [16]. A large majority of them were also absent from the model strain B05.10 [42] and were predicted without RNAseq data. In order to better characterize these Vv3 accessory genes and investigate whether they could be cargo genes carried by DNA transposons, a new prediction and annotation was conducted based on the RNAseq data generated in this study (See Methods). A total of 281 genes were identified as present in the genome of Vv3 but not in the genome of Sl3 (Table S6). Thirty-seven percent of these Vv3 genes (n = 104) were located on the AC19, the majority of which (77) were either not expressed or exhibited very low expression levels. When considering the most expressed ones, only few functions could be predicted including mainly peptidases (3 genes) and activities in relation with DNA and RNA regulation (nucleases, helicase, transcription factors).

The other Vv3-specific genes (177 genes) were localized in the core chromosomes, mainly in the subtelomeric regions as previously observed [16]. In particular, a set of 45 Vv3-specific genes (Vv3_03_G_03414-3458) were co-localized in a 93Kb subtelomeric region of the chromosome 3 (Fig. 4) that is absent from the genomes of the Sl3 and B05.10 strains. At one end of this region, the newly provided annotation identified two expressed genes (Vv3_03_G_03450 and 03458) that encode proteins with a DUF3435 domain which is a specific feature of the giant transposons called *Starships* [7]. This protein, encoded by the first gene of every *Starship*, is a tyrosine recombinase (YR) that mediates the movement of the TE and has been therefore named the “captain” [6]. In addition to these two captain genes, the Vv3 region carries 43 cargo genes including four genes with functional predictions that are the most commonly found across *Starships i.e.* a DUF3723-protein, a ferric reductase (FRE), a patatin-like phosphatase (PLP) and a Nod-like receptor (NLR) [7]. Finally, as in several other *Starships*, the gene at the opposite side of the 5’ captain gene encodes a SANT/Myb transcription factor [57]. Two other key features define *Starships*: the flanking direct repeats (DRs) that are generated by the insertion of DNA transposons, and the terminal inverted repeats (TIRs) required for the excision [7, 9, 12]. In the case of the Vv3 genome, the insertion of the *Starship* occurred in a low complexity AT-rich region composed of numerous embedded TEs which impaired the identification of TIRs and DRs. Nevertheless, the annotated genes indicated that the identified 93 kb Vv3-specific region is a *Starship* element. To classify this new TE, we compared the sequence of both captain Vv3 proteins with the HMM profiles of YR reference sequences, as recommended by Gluck-Thaler and Vogan [9]. The HMM profiles of the two Vv3 captain proteins revealed high similarities with the captains of the *Phoenix* family of *Starships*. Then a phylogenetic analysis confirmed the position of the Vv3 *Starship* among the *Phoenix* family (Fig. S7; Table S7). A blast analysis against the Starbase database dedicated to *Starships* [58] failed to identify homology with a previously identified *Starship*. The newly identified *Starship* of the Vv3 strain was named *Ariane* after the name of the European space rocket whose name start with the same letters than arsenic (see below), but also in reference to the Greek myth of *Ariadne* who married *Dionysus*, the god of wine.

**Fig. 4.**
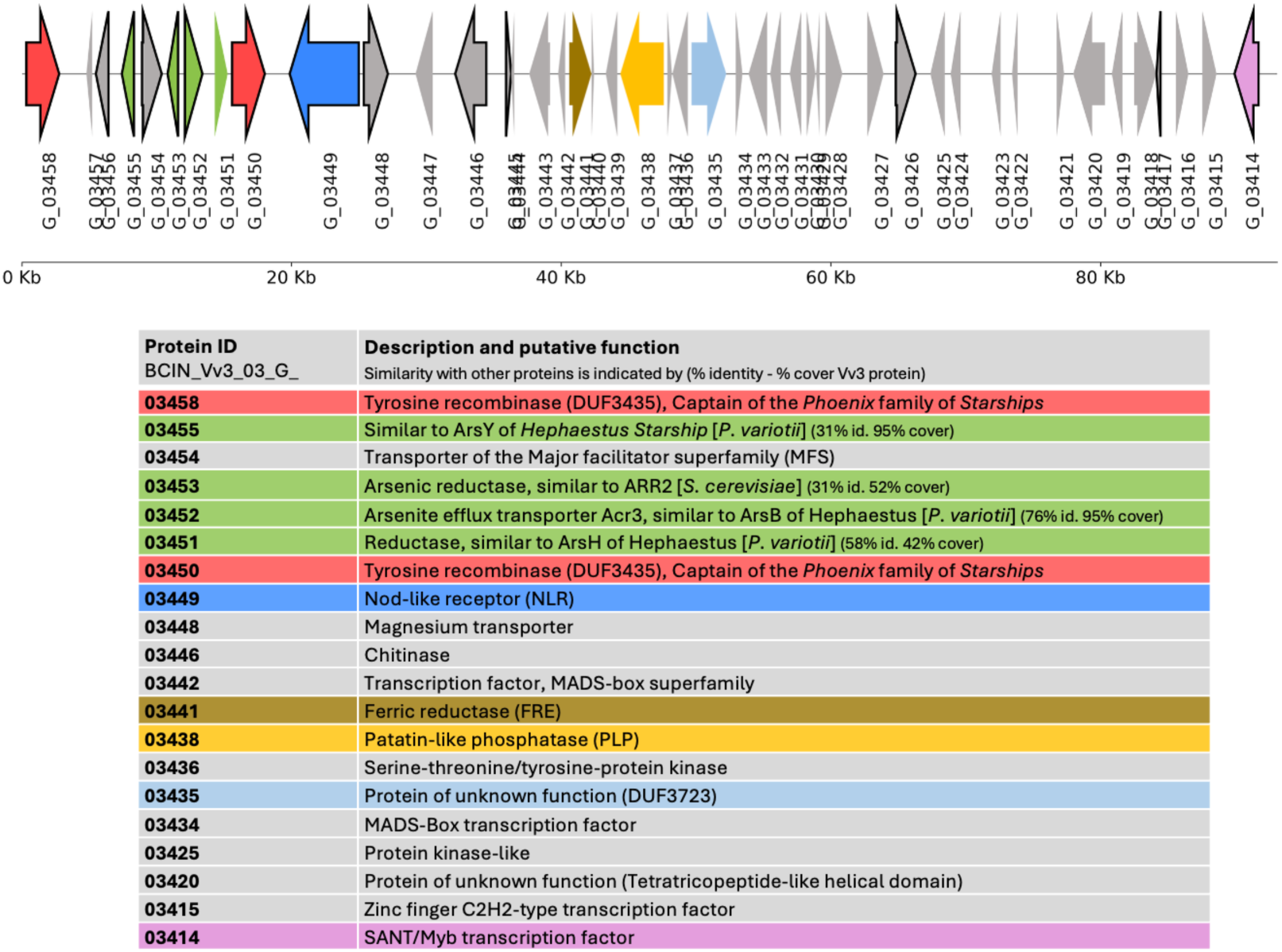
A *Starship*, called *Ariane,* was identified on the chromosome 3 of the *B. cinerea* strain Vv3 (BCIN_Vv3_03: 3,353,355-3,446,435). 45 genes were annotated within this 93 Kb-transposon. Outlined genes showed evidence of expression at 24h or 48h post-inoculation on grapevine berries (normalized counts > 50 in at least one condition). The table lists the genes with predicted functions. Typical proteins of *Starships* are highlighted in colors. Two genes were predicted to encode tyrosine recombinases, *i.e.,* the captains of *Ariane* (in red). Based on the phylogeny of these two proteins, *Ariane* was classified as a member of the *Phoenix* family of *Starships*.

### *Ariane* is present in additional strains of *B. cinerea* specialized to grapevine

The presence of *Ariane* in fungi was further investigated by BLASTn against all the genomes available at NCBI. In addition, we used an alternative approach based on protein similarity *i.e.,* searching clusters of co-located homologous sequences against nr NCBI using cblaster [59]. In both analysis, *B. cinerea* was the only organism in which *Ariane* was detected. The distribution of *Ariane* was then investigated in the 64 available genomes of *B. cinerea* strains isolated from grapevine (22 strains), tomato (15 strains), strawberries (5 strains), petunia (4 strains) and various other hosts. BLAST analysis indicated that *Ariane* was present either in complete or incomplete copies in 18 strains, mostly isolated from grapevine (Table S1). When considering the copies with a size closely related to those of the Vv3 copy (>80 kb), only five strains were identified, four of them belonging to the French populations specialized to grapevine (Vv3, Vv8, Vv12 and Vv14; [14]) and one isolated from the same host in Spain (Bc448) (Fig. S8). In contrast, no trace of *Ariane* could be detected in the 15 genomes of Sl strains specialized to tomato [14].

### *Ariane* confers resistance to arsenic

In addition to the genes with functional predictions commonly found across *Starships* i.e. those mentioned above, *Ariane* was found to carry genes encoding proteins with various IPR domains highlighting several transcriptional regulators, transporters and a chitinase (Table S6; Fig. 4). Some of these genes didn’t show evidence of expression in the tested conditions (infection of grape berries) and could correspond to pseudogenes. Nevertheless, the region localized between the two captain genes retained our attention as two genes with evidence of expression were predicted to encode proteins whose IPR domains suggest a possible role in arsenic resistance (Table S6): BCIN_Vv3_03_G_03452 carries the IPR domain of the yeast arsenite efflux transporter Acr3 [60] and BCIN_Vv3_03_G_03453 carries a Rhodanese-like domain like the yeast arsenic reductase ARR2 [61]. A *Starship* (*e.g. Hephaestus*) identified in the soil fungus *Paecilomyces variotii* was shown to contain a subregion involved in arsenic resistance [11]. Predicted proteins encoded by genes of *Ariane* and those of *Hephaestus* were therefore compared. The efflux transporter encoded by BCIN_Vv3_03_G_03452 showed 76% identity with the ArsB protein of *Hephaestus.* In addition, the proteins encoded by the neighbor genes BCIN_Vv3_03_G_03451 and BCIN_Vv3_03_G_03455 showed similarities with the putative arsenate reductase ArsH and the protein of unknown function ArsY of *Hephaestus* [11]. While *Hephaestus* and *Ariane* were found to share three genes (*ArsY, ArsB* and *ArsH*), no synteny was observed between them.

As four cargo genes of *Ariane* could be related to arsenic resistance, experiments were designed to test this hypothesis in *B. cinerea.* Conidial germination and hyphal development were observed in presence of different concentrations of sodium (meta)arsenic AsNaO_2._ Conidia suspended in PDB were inoculated on hydrophobic coverslips. At 20 hours post-inoculation, the formation of successive appressoria-like structures at the tips of the swollen hyphae could be observed for both Vv3 and Sl3 strains (Fig. 5., top pictures). At 750 μg/mL AsNaO_2_, no germination was visible for Sl3. In contrast, 98.5% of the conidia of the Vv3 strain germinated although they showed a delay in hyphal elongation (Fig. 5., bottom pictures). At 3 days post-inoculation, Sl3 conidia were still ungerminated while mycelium was fully developed for the Vv3 strain indicating a resistance to arsenic (data not shown). The resistance of the grapevine specialized strains Vv1 to Vv16 was tested in similar conditions and only the strains carrying the *Ariane* element (Vv8, 12, 14) or at least the 5’ part including the genes with predicted functions related to arsenic resistance (Vv16; Fig S8) were able to germinate and develop mycelium in the presence of 750 μg/mL AsNaO_2_.

**Fig. 5.**
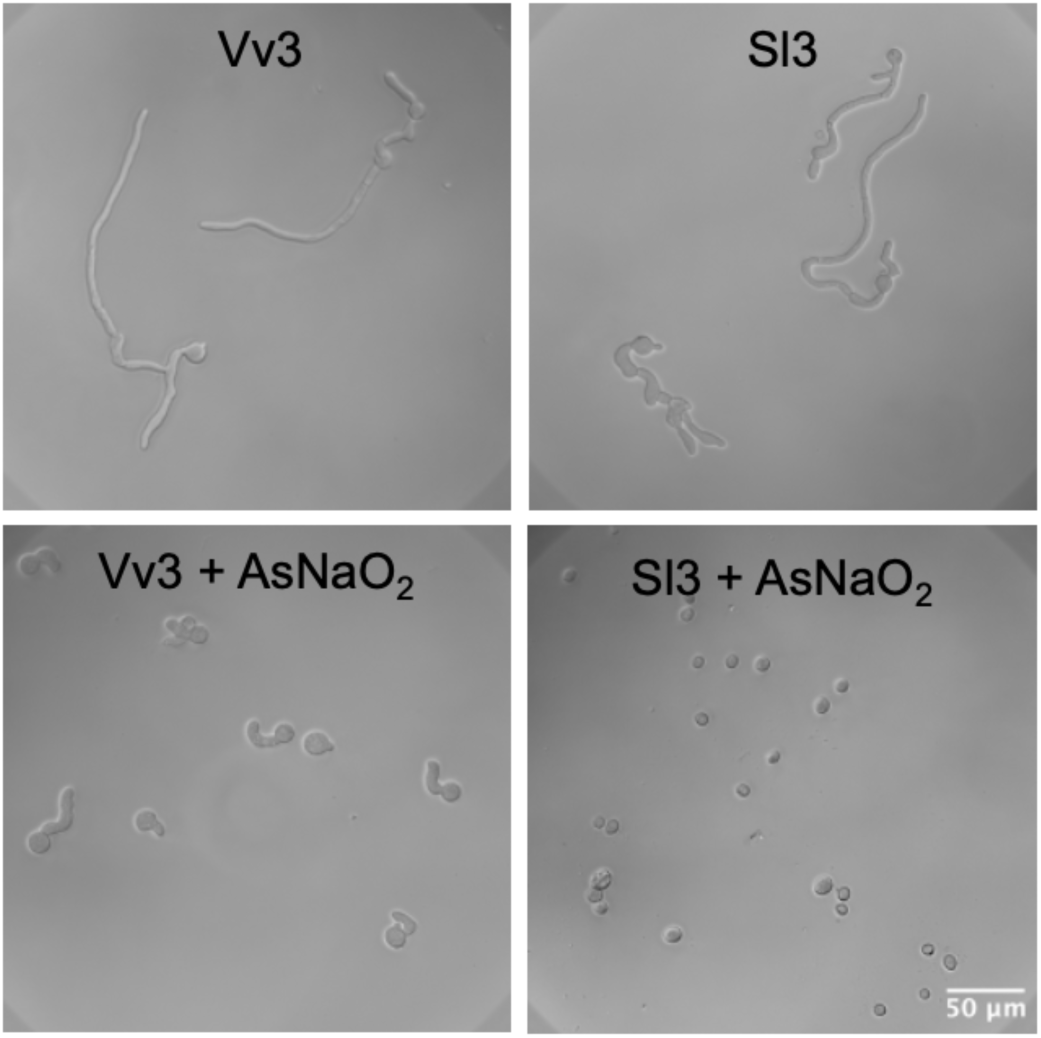
Germination of *Botrytis cinerea* conidia on hydrophobic surface and inhibitory effect of arsenic. Conidia of the strains Vv3 and Sl3 were inoculated in PDB medium diluted to the fourth and incubated on glass coverslips for 20h at 12°C. In the bottom pictures, 750 μg/mL AsNaO_2_ were added to the medium.

To establish whether arsenic resistance was due to the presence of *Ariane*, a formal genetics approach was used. A cross between the Vv3 and Sl3 strains was performed and 100 progenies were analyzed for their resistance to arsenic (as described above) and for the presence of *Ariane*. PCR primers were designed to detect the captain genes, three genes possibly involved in arsenic resistance (BCIN_Vv3_03_G_03452, G_03253 and G_03454) and the gene at the 3’ end of the *Starship* encoding a SANT/Myb transcription factor (G_03414). Primers were also designed to detect additional loci specific of either the parental strain Vv3 (*BcMat1-1* on chromosome 1, *BcMelA* on chromosome 2) or Sl3 (*BcMat1-2* on chromosome 1, *BcStc6* on chromosome 3, [16]). As expected, the progenies showed recombination between the parental markers (Table S8). Segregation of the *BcStc6* genes and genes of *Ariane* further confirmed that intrachromosomal recombination occurred in chromosome 3 as the progeny were either of the parental genotype (with the *BcStc6* gene or with *Ariane*), either of a recombinant genotype (with both *BcStc6* and *Ariane* or without any of them). In two cases (progenies #20 and #23), only the SANT/Myb encoding gene was detected suggesting that only a 3’ part of *Ariane* was inherited. Finally, a strict correlation was identified between the presence of all five *Ariane* PCR markers and growth in presence of arsenic, indicating that this *Starship* confers the observed resistance (The probability of obtaining such data under the assumption of independent traits is less than 10⁻²³).

## DISCUSSION

The polyphagous pathogen *Botrytis cinerea,* owing to the existence of host-specialized populations [14, 15, 62] and to a specific repertoire of transposons in the strains specialized on grapevine [16], provides a particularly relevant model to investigate the role of TEs in adaptation. In this study, both the role of retrotransposons and that of a giant DNA transposon were investigated. The results indicate that retrotransposons-derived small RNAs do not have a significant role in the virulence of *B. cinerea* on grapevine, but revealed a so far unknown *Starship* whose cargo genes provides resistance to arsenic and could therefore contribute to the adaptation of strains to cultivated grapevine.

Several population studies demonstrated that the repertoires of retrotransposons of the *Copia* and *Gypsy* families are highly strain-dependent in *B. cinerea*. Some of these TEs are widely distributed and were detected in strains isolated from diverse hosts. That’s the case of *BcGypsy1* (also named *Boty*)*, BcGypsy3* and *BcGypsy4* all known to be the source of small RNAs including *Bc*siR5, *Bc*siR3.1 and *Bc*siR3.2 that act as effectors targeting the transcripts of conserved host defense genes [4, 63]. In opposite, three sRNA generating TEs have only been identified in grapevine specialized strains so far i.e. *BcCopia4, BcGypsy6* and *BcGypsy7* which could suggest that they specifically target *V. vinifera* genes [16]. In order to test the role of retrotransposon-derived RNA in the interaction with grapevine, Dicer mutants unable to produce sRNA were generated in the Vv3 strain. In the model strain B05.10, inactivation of both *BcDcl1* and *BcDcl2* genes was required to abolish siRNA production, indicating a redundant role of the two BcDCL proteins in the production of sRNAs [4]. In Vv3, the inactivation of *BcDcl1*^Vv3^ was sufficient to abolish the production of *Bc*siR3.1, *Bc*siR3.2 and *Bc*siR5 and the Vv3-specific *Bc*siR1^Vv3^, suggesting that in this strain *BcDcl2*^Vv3^ could not take over the generation of these sRNA. This difference could not be explained by a difference in gene sequence (100% identical between Vv3 and B05.10) nor by a lack of expression. In the B05.10 strain, the *ΔΔBcdcl1/2* double mutant unable to produce DCL-dependent sRNAs was first reported to generate smaller lesions than the wild-type strain on plant tissues *i.e., Arabidopsis thaliana* and tomato leaves. In addition, this mutant displayed a growth defect and sporulation delay [4]. However, the same B05.10 *ΔΔBcdcl1/2* double mutant made in an independent study was fully virulent [64]. Our data obtained with the Vv3 grapevine-specialized strain are consistent with this second study as the *ΔBcdcl1^Vv3^* and *ΔΔBcdcl1/2^Vv3^* mutants didn’t show any significant pathogenicity defect on grapevine berries or leaves. This indicates that DCL-dependent sRNAs are not required for the necrotrophic development on grapevine. Nevertheless, this doesn’t mean that CK RNAi doesn’t occur between *B. cinerea* and grapevine. The putative target grapevine genes of the small RNA generated by the retrotransposons *BcCopia4, BcGypsy6* and *BcGypsy7* were searched by combining *in silico* prediction of the sRNA-target mRNA duplexes and transcriptomics. Twenty-one candidate genes were identified. These 21 host genes are less expressed when grapevine berries are infected by the strain Vv3 than during infection with the strain Sl3 (that lacks the *BcCopia4, BcGypsy6* and *BcGypsy7* TEs) which is compatible with a silencing effect. To our knowledge, no study conducted with *B. cinerea* brought the proof of the occurrence of expected host target mRNA cleavage in the host plant. In our study, we set up a RLM-RACE experiment [31] to verify that the 21 candidate host mRNAs were cleaved at the predicted sites. Unfortunately, while we experimentally identified cleavage sites on the candidate mRNA, those were different from the predicted ones and didn’t confirm the action of the fungal sRNA. At this stage, it is difficult to conclude about the occurrence or not of the cleavage at the predicted sites. Indeed, the lack of evidence could also be due to technical limitations or secondary cleavages. In conclusion, this first study designed to investigated sRNA produced by a *B. cinerea* strain specialized on grapevine didn’t provide any evidence of a significant role of these secreted effectors on the virulence toward the host of origin. Unlike previous studies, our data are based on a strain whose original host and host specificity are known, which should yield results that more accurately reflect what occurs under natural conditions. While no impact on virulence could be detected in our pathogenicity tests, it remains possible that these sRNA play a role in the infection of berries at early stage of their development. Indeed, in the vineyards, *B. cinerea* can infect senescent tissues of flowers and enter young berries at fruit set stage, then start a latency phase during which the pathogen is quiescent without causing disease symptoms, generally until berries begin to ripen [23, 65]. It would be interesting to investigate whether *B. cinerea* sRNA act on grapevine immunity to prevent defense reactions during this latent stage.

Exploring the accessory genes present in the Vv3 genome, our study revealed a new 93 kb-*Starship* transposon that we named *Ariane*. Despite their large size and numerous cargo genes, *Starships* were long hidden in fungal genomes. Thanks to the recent characterization of their specific features especially the captain gene that encodes the tyrosine recombinase (YR) responsible for their transposition, they are now being identified in many fungal genomes [8, 9]. *Ariane* contains two captain genes encoding the same YR whose HMM profile and phylogeny indicated a *Starship* of the *Phoenix* family. The presence of two captain genes could be the result of an imbrication of two elements. *Starships* can indeed sequentially nest within each other and complex rearrangements have been described in several species [7, 11, 57]. In addition to the two captain genes, *Ariane* contains several cargo genes that are typical features of *Starships* i.e. those encoding a DUF3723-protein, a ferric reductase (FRE), a patatin-like phosphatase (PLP) and a Nod-like receptor (NLR). While the exact functions of these genes are not known, one hypothesis is that they could be involved in the mobilization of host genes and in horizontal gene transfers (HGTs). Investigating hundreds of fungal genomes, Gluck-Thaler et al. [9] revealed that *Starships* would indeed be responsible for many HGTs between distantly related Ascomycetes and would play a major role in the supply of accessory genes. *Starships* are therefore considered to have a major role in fungal adaptation and in fungal disease outbreaks. One striking example is the case of the necrotrophic effector ToxA carried by different *Starships* across unrelated fungal plant pathogens [12]. The *Ariane* element identified in the Vv3 strain and in other strains all isolated from grapevine was found to significantly contribute to the accessory compartment of the genome. The 45 genes within this giant TE correspond to 16% of the 281 genes that are present in the Vv3 genome but not in the Sl3 genome. Nevertheless, some of the 45 predicted genes were short in size and didn’t show any expression which could be the sign of pseudogenization like in other *Starships* [11]. Among the cargo genes of *Ariane* with a significant level of expression, several ones showed predicted functions that could contribute to fungal adaptation to specific environments. In particular, genes encoding proteins related to arsenic resistance were identified in 5’ extremity of *Arian*e. Resistance to arsenic was therefore investigated by testing the germination of conidia in the presence of AsNaO_2_. While the Sl3 strain was completely inhibited at 750 μg/mL AsNaO_2_, all Vv strains carrying *Ariane* (Vv3, 8, 12, 14 and 16) germinated indicating resistance to this toxic metalloid. Finally, a genetic cross between the strains Vv3 and Sl3 demonstrated that *Ariane* confers resistance to arsenic as only the half of progenies that carries *Ariane* is resistant to arsenic. Notably, the *Hephaestus Starship*, named after the Greek god of metallurgy, was recently found to confers protection from toxic levels of different metal ions including arsenic in the fungus *P. variotii* [11]. On the basis on their captain sequence, *Ariane* and *Hephaestus* belong to different families of *Starships* but they share three cargo genes *i.e., ArsH*, *ArsB* and *ArsY*. In *P. variotii,* the subregion of *Hephaestus* that contains six *Ars* genes was also experimentally shown to be responsible for resistance to arsenic. In both *Hephaestus* and *Ariane*, the multiple *Ars* genes could act in synergy to confer resistance to arsenic. Therefore, several simple and multiple gene deletions would be required to evaluate the contribution of each of them. Beside *Starships* in filamentous fungi, genes involved in arsenic resistance were already shown to be organized into clusters in *Saccharomyces cerevisiae* [66] and in operons in procaryotes [67]. These clusters or operons combine genes involved in efflux and detoxification of arsenic. In the case of *Ariane*, the candidate genes involved in arsenic resistance could be responsible for arsenic reduction (proteins similar to Arr2 and ArsH), or efflux (protein similar to both Acr3 and ArsB; Fig. 4). Additional cargo genes in the *Ars* region of *Ariane* could be involved in the phenotype especially the one encoding transporter of the major facilitator superfamily (MFS; BCIN_Vv3_03_G_3454) that could allow the efflux of toxic compounds. Among the 64 *B. cinerea* strains whose genomes are available at NCBI, 18 were found to carry *Ariane* or remaining traces of it. Complete copies of *Ariane* were found in a sub-selection of five strains, all isolated from grapevine in different French regions or in Spain. As genetic exchanges occur between populations of *B. cinerea* specialized on different hosts [14, 15], the detection of complete *Ariane* copies only in grapevine isolated strains so far suggests that this *Starship* confers an ecological advantage on this host or its environment. Actually, sodium arsenite was widely used in viticulture during the 20^th^ century to fight against insects and fungi causing trunk diseases such as Esca. Using a molecular metabarcoding approach, a study recently confirmed that such treatment can significantly reduce the abundance of the Esca-associated fungus *Fomitiporia mediterranea* but can also significantly impact the mycobiota colonizing grapevine woody trunk tissues [68]. Due to its toxic effect on a wide range of organisms, arsenic was banned in 2001 in France and 2003 in Europe [69]. The Vv strains investigated in our study were collected in France in 2006 when traces of the metalloid were still present in the soil and in grapevine organs [70] and were probably exerting selection pressure on *Ariane* in *B. cinerea*. Wider pangenomic studies in *B. cinerea* would further allow to confirm the link between sodium arsenite treatments and the occurrence of *Ariane.* The functional role and possible selective advantages provided by the other cargo genes also remain to be investigated. Investigating the presence of *Ariane* or related *Starships* in other fungal species, in particular those present in the grapevine mycobiota, may help to understand their evolutionary histories and to detect possible HGTs. More generally, in a context where fungal pathogens are becoming increasingly resistant to antifungal treatments in both agriculture and medicine [71], the capacity of *Starships* to transfer dozens of genes through between taxonomically distant fungi HGTs [9, 72] should be fully considered in a One Health approach. In conclusion, this study highlights how a *Starship* giant transposon shaped the accessory genes compartment of the polyphagous fungus *B. cinerea* and contributed to its adaptation to vineyards by conferring resistance to arsenic.

## Acknowledgements

We are grateful to the following bioinformatics platforms and partners for providing computational support and/or storage resources: Bioinfo Genotoul, (https://doi.org/10.15454/1.5572369328961167E12), CATI BARIC (https://www.cesgo.org/catibaric/) and Nicolas Lapalu (INRAE-BIOGER Bioinformatics platform). We also thank Hailing Jin and Baoye He (University of California, USA) for sharing the anti-AGO1 antibodies as well as Mathilde Faguard (INRAE Versailles, France) for hosting A. Porquier in her lab during the move of the BIOGER unit.

## Author contributions

A.P. Conceptualization, supervision, experiments, formal analysis, writing; A.S. Conceptualization, data curation, formal analysis, writing, editing; J.Ve. Experiments, formal analysis; J.Vi. Experiments, formal analysis, S. A. Experiments, S.B. Supervision, writing, J.C. Experiments; A.C. Experiments; J.R. Methodology; A. D. Methodology; A.S.W. Resources, editing; M.A. Funding acquisition, writing, editing; B.P. Funding acquisition, conceptualization, editing; M.V. Funding acquisition, conceptualization, supervision, experiments, formal analysis, writing and editing.

