## Supplementary figures for "Role of transposons in the specialization of *Botrytis* cinerea to grapevine: Insights into small RNAs and a new *Starship*"

Key words: Retrotransposons, Small RNAs, DICER proteins, *Starship*, Arsenic resistance

Antoine Porquier[1], Adeline Simon[1], Justine Vergne[1], Jérémy Villette[2], Sébastien Aimé[2], Stéphane Bourque[2], Jérôme Chapuis[1], Anthony Colas[1], Justine Rouffet[1], Antoine Davière[3], Anne-Sophie Walker[1], Marielle Adrian[2], Benoit Poinssot[2], Muriel Viaud[1 *]

[1] Université Paris-Saclay, INRAE, Biology of fungal plant pathogens (BIOGER), Palaiseau, France

[2] Agroécologie, Institut Agro Dijon, INRAE, Université Bourgogne Europe, Dijon, France

[3] Université Paris-Saclay, INRAE, AgroParisTech, Institute Jean-Pierre Bourgin for Plant Sciences (IJPB), 78000, Versailles, France

[*] Corresponding author

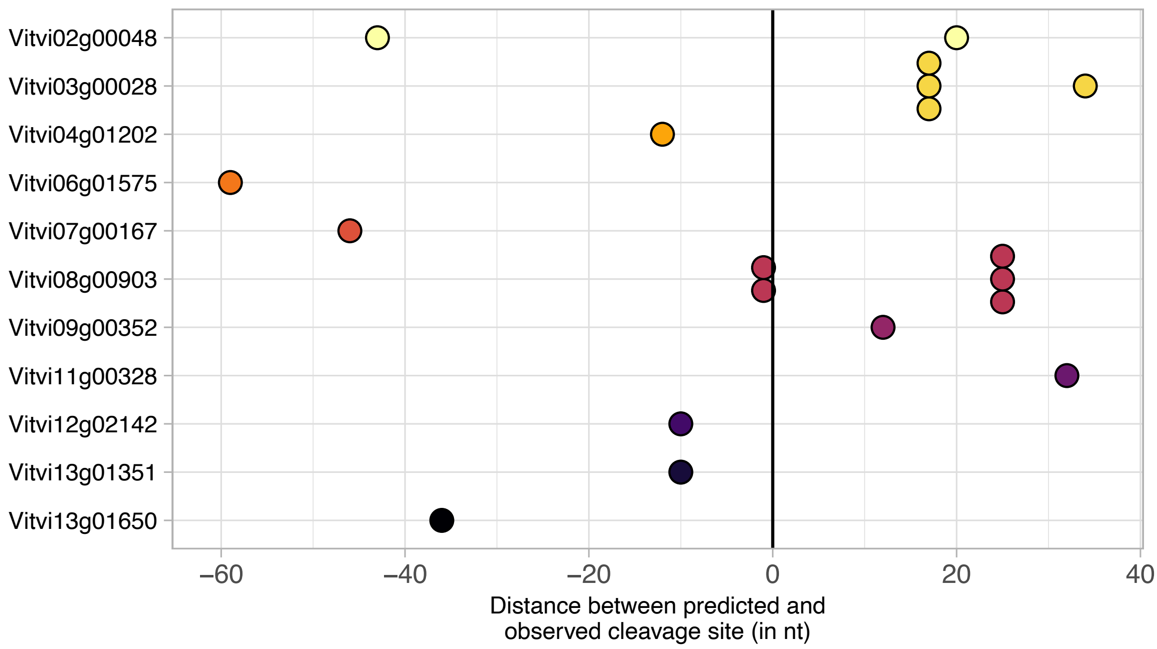


**Fig. S1**. Identification of the cleavage sites of grapevine mRNAs during infection by the *Botrytis cinerea* strain Vv3. Grapevine mRNAs predicted to be targeted by fungal small RNAs were amplified by RLM-RACE PCR. One to five PCR products by mRNAs were sequenced. The sequences were aligned on the corresponding reference target mRNAs and the distance (in nt) between the predicted cleavage sites and the observed ones were plotted: a negative value indicates a cleavage upstream while a positive score indicates an observed cleavage downstream of the expected site.

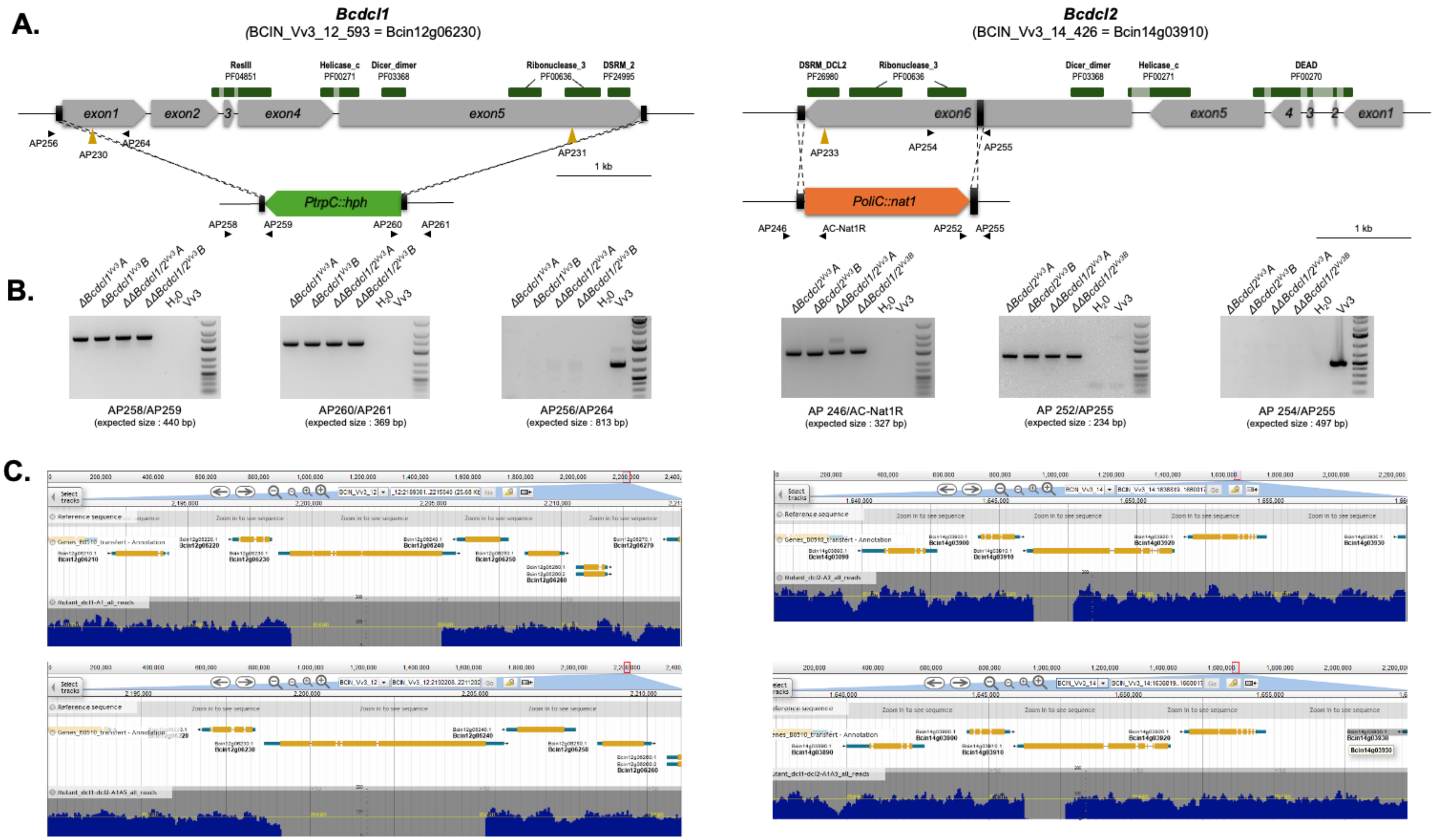


**Fig. S2.** Inactivation of the *BcDcl1* and *BcDcl2* genes in the *Botrytis cinerea* strain Vv3. A. Schematic view of the genes, protein domains, PCR primers, guide RNAs (in brown) and deleted region. B. PCR results indicating the integration of the *hph* and *nat* gene at the expected insertion site and the absence of WT copies of the gene in the mutants. C. Validation of the deleted regions by full genome sequencing.

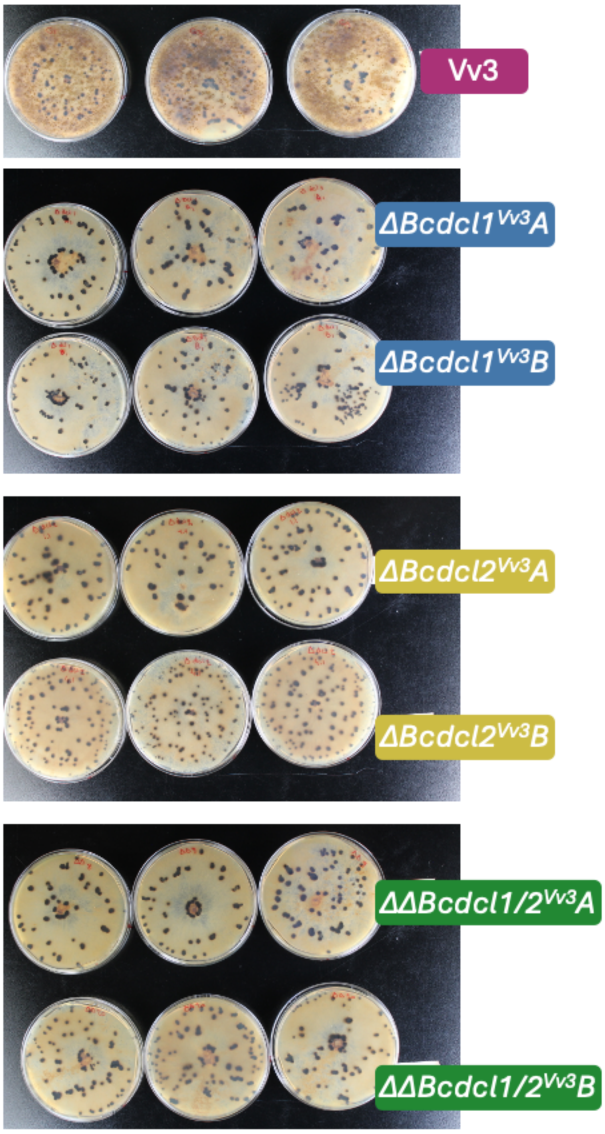


**Fig. S3.** Sclerotia production in *Botrytis cinerea* strains. The ability to produce sclerotia was assessed by growing the strains at 4°C in the dark for one month. Three technical replicates were realized and are presented here.

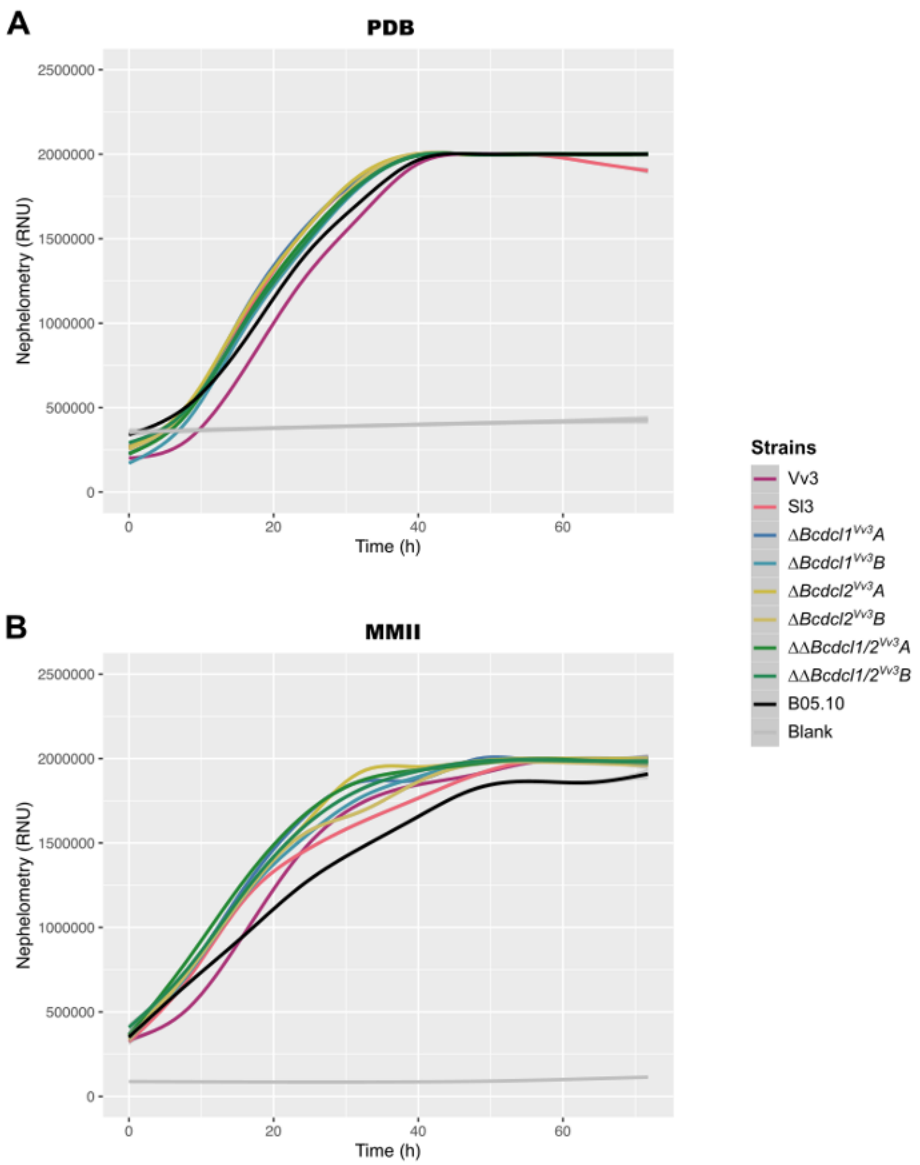


**Fig. S4.** Growth curves of the different strains of *Botrytis cinerea* compared by nephelometry. Conidial suspensions were prepared at a concentration of $10^4$ conidia/mL in riche (PDB) or minimal medium (MM) and cultivated for 72 hours at 20° C in 96-wells plates. Fungal growth was recorded by using the NEPHELOstar Plus device. The results plotted here were obtained from raw results of 5 technical replicates (3 for the blanks). Growth is measured in relative nephelometric units (RNU).

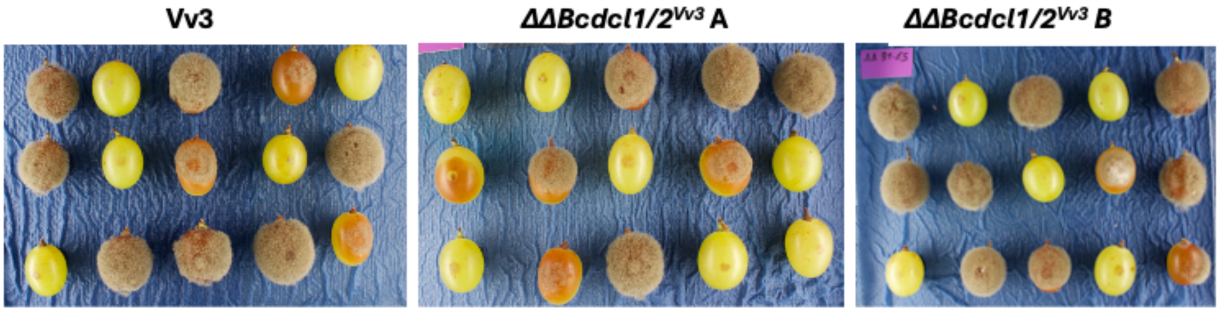


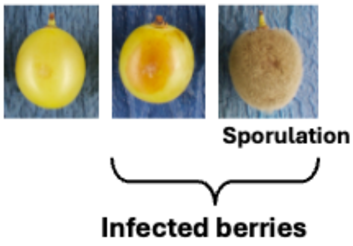


| Strain | Infected berries | Infected berries with sporulation |
|---|---|---|
| WT Vv3 | 70% (±19) | 64% (±16) |
| *ΔΔBcdcl1/2*$^{Vv3}$ A | 72% (±17) | 61% (±22) |
| *ΔΔBcdcl1/2*$^{Vv3}$ B | 82% (±17) | 69% (±11) |
| WT Sl3 | 54% (±20) | 45% (±26) |

**Fig. S5.** Virulence assays of *Botrytis cinerea* strains on grapevine berries. Berries were inoculated with 1000 conidia resuspended in potato dextrose broth. The photographs show representative results obtained with the WT strain Vv3 and the two *ΔΔBcdcl1/2* $^{Vv3}$ double mutants at 11 dpi. The table indicates the average percentages of infected berries and of those with sporulation from three experiments with 15 to 22 berries per strain.

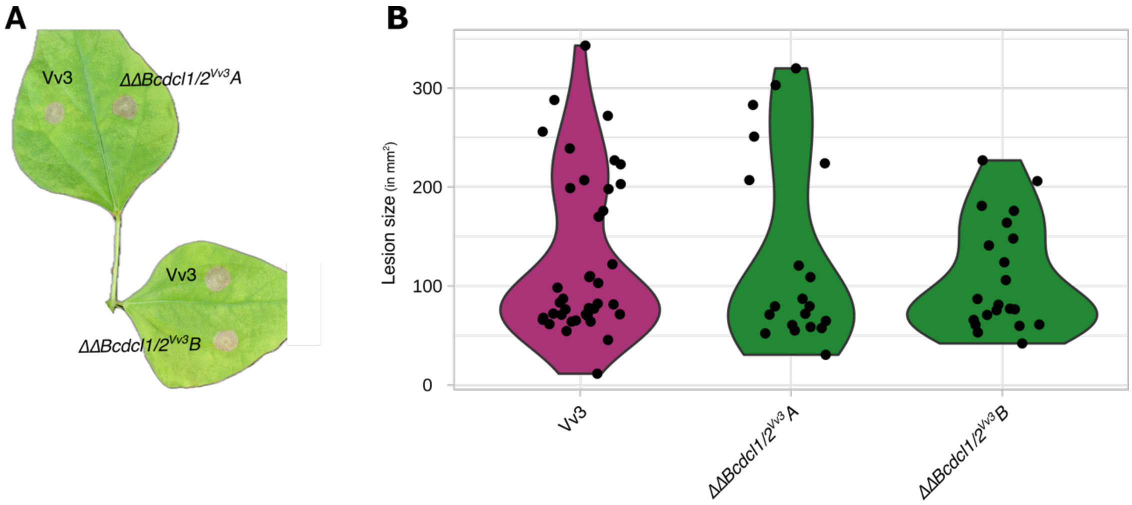


**Fig. S6.** Virulence assays of *Botrytis cinerea* strains on French bean. Leaves were inoculated with 1000 conidia and incubated four days. A. Representative results obtained with the WT strain Vv3 and the *ΔΔBcdcl1/2* $^{Vv3}$ double mutants. B. Average lesion size of the symptoms calculated from two independent biological replicates with at least 9 inoculations (technical replicates) for each modality. Statistical analysis (Kruskal-Wallis) showed no significant differences between the WT and the mutants.

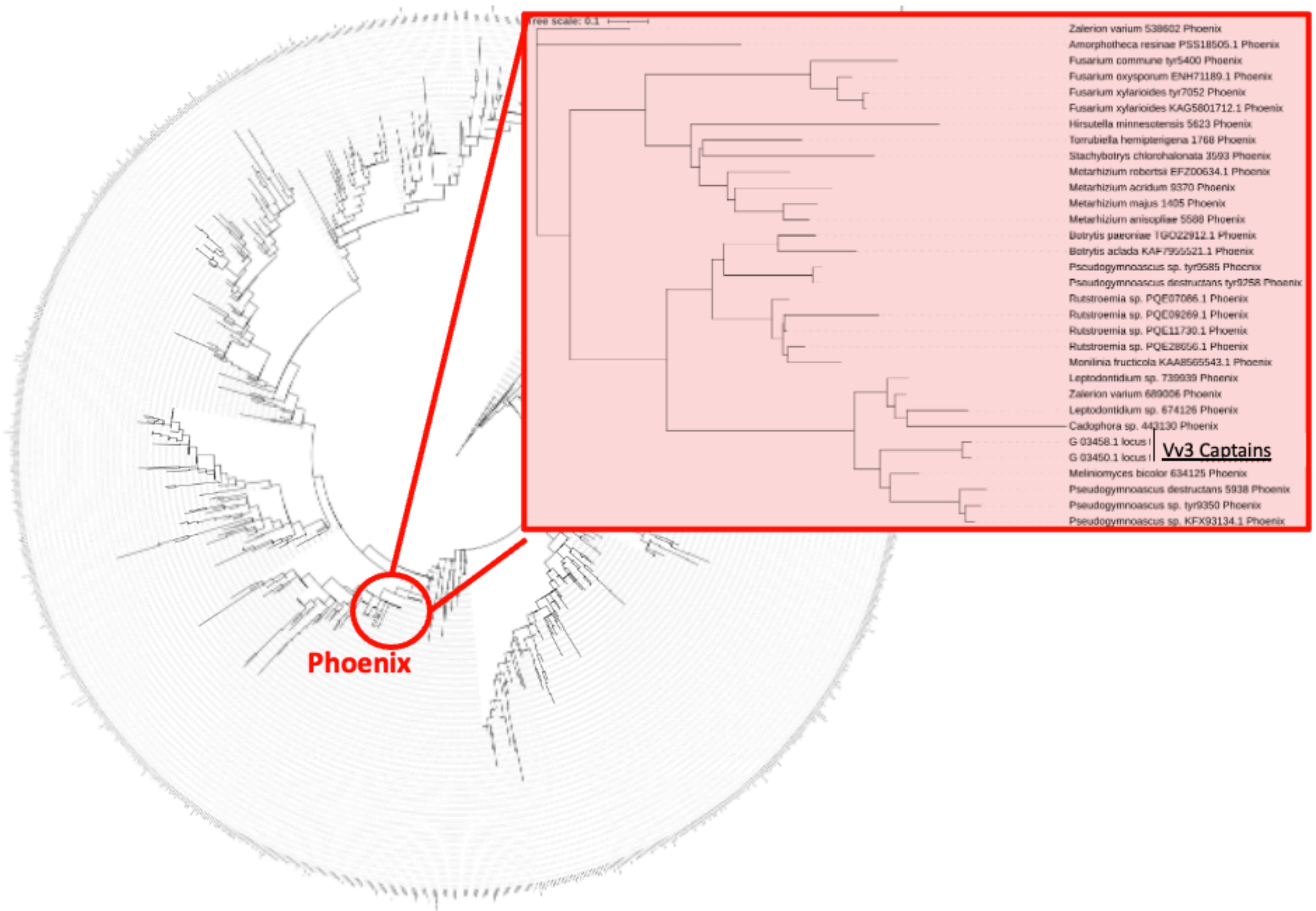


**Fig. S7.** Phylogenetic position of the tyrosine recombinases (YRs) encoded by genes of the *Ariane* TE identified in the *Botrytis cinerea* Vv3 strain indicating that this *Starship* belongs to the *Phoenix* family. An evolutionary tree of 1222 representative *Starship* YRs and the two Vv3 YR was built from an alignment of protein sequences. The box shows the tree derived from the alignment of the sequences from the red node, that includes the sequences of the two Vv3 YRs.

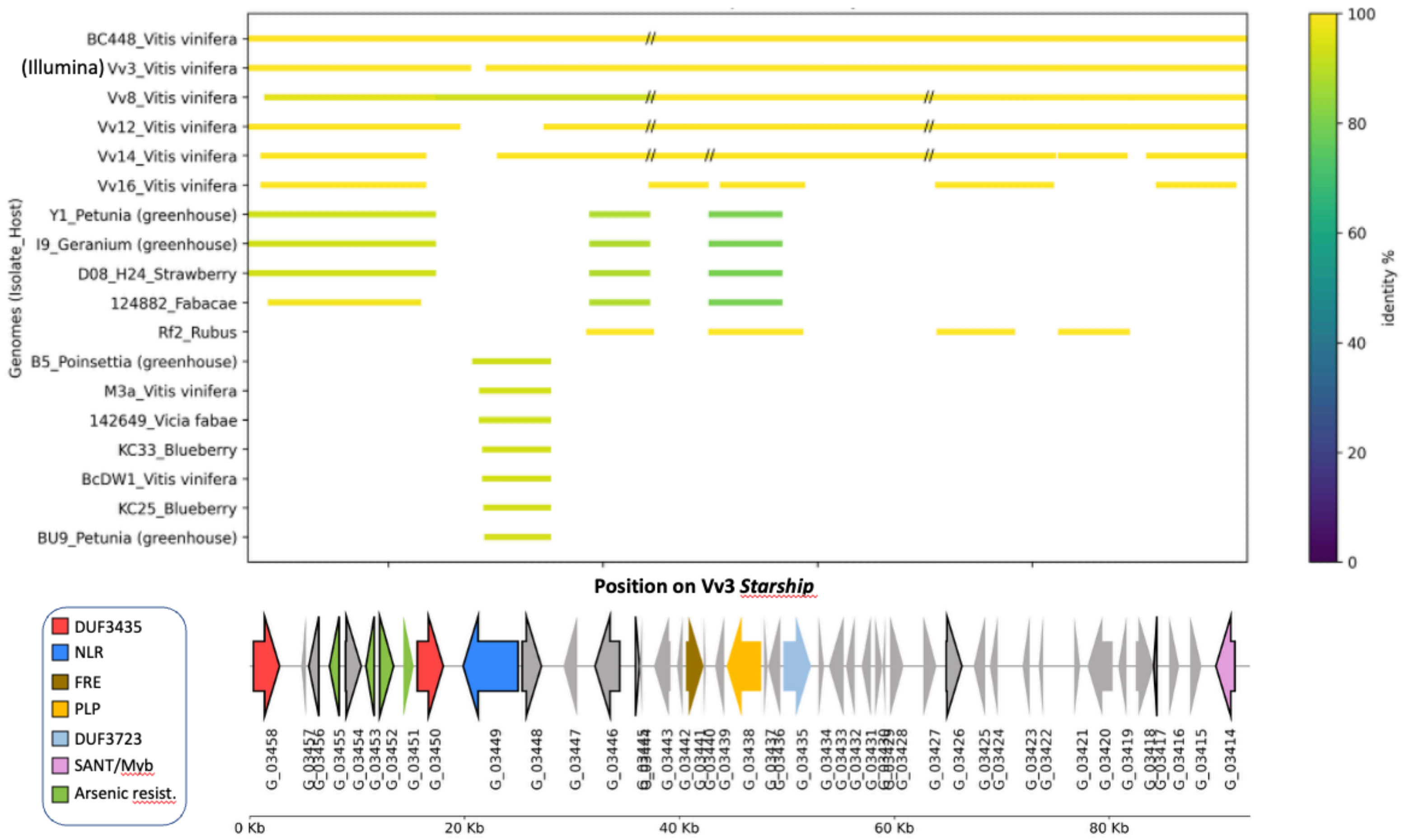


**Fig. S8**. Distribution of *Ariane* in *Botrytis cinerea* strains from different hosts and origins. All 64 *B. cinerea* genomes available at NCBI (https://www.ncbi.nlm.nih.gov/datasets/genome; Table S1) were queried against the sequence of *Ariane* identified in the Vv3 strain. Only blast alignments with a length > 5Kb were displayed.
